# Morphological determinants of bite force capacity in insects: a biomechanical analysis of polymorphous leaf-cutter ants

**DOI:** 10.1101/2021.08.06.453764

**Authors:** Frederik Püffel, Anaya Pouget, Xinyue Liu, Marcus Zuber, Thomas van de Kamp, Flavio Roces, David Labonte

## Abstract

The extraordinary success of social insects is partially based on ‘division of labour’, i. e. individuals exclusively or preferentially perform specific tasks. Task-preference may correlate with morphological adaptations so implying task-specialisation, but the extent of such specialisation can be difficult to determine. Here, we demonstrate how the physical foundation of some tasks can be leveraged to quantitatively link morphology and performance. We study the allometry of bite force capacity in *Atta vollenweideri* leaf-cutter ants, polymorphic insects in which the mechanical processing of plant material is a key aspect of the behavioural portfolio. Through a morphometric analysis of tomographic scans, we show that the bite force capacity of the heaviest colony workers is twice as large as predicted by isometry. This disproportionate ‘boost’ is predominantly achieved through increased investment in muscle volume; geometrical parameters such as mechanical advantage, fibre length or pennation angle are likely constrained by the need to maintain a constant mandibular opening range. We analyse this preference for an increase in size-specific muscle volume and the adaptations in internal and external head anatomy required to accommodate it with simple geometric and physical models, so providing a quantitative understanding of the functional anatomy of the musculoskeletal bite apparatus in insects.

## Introduction

Colonies of social insects are stereotypically characterised by a ‘division of labour’, i. e. individuals favour some tasks over others [1, 2]. Such task-preferences may be the ‘plastic’ result of complex interactions between genetic predisposition, neural and hormonal factors, and varying levels of experience [2, 3], but they may also correlate with more ‘rigid’ morphological differences in the colony workforce [4, 5]. A natural aim, then, is to connect morphological variation with the task-specialisation it enables.

A textbook example of polymorphic social insects are leafcutter ants, in which workers fall on a more or less continuos size-spectrum covering more than two orders of magnitude in body mass (see Fig. 1). Workers may not only differ in size, but also in shape, i. e. specific body parts may not scale according to geometric similarity hypotheses [6–13]. Both size- and shape-variation may indicate task-specialisation and so help to increase colony fitness through ergonomic task allocation [7, 10, 14–20], but this link is speculative where it is not rationalised through a quantitative functional analysis. For some tasks, e. g. brood care or gardening, such an analysis may be difficult to conduct, but there exists a subset of tasks where it can be made both explicit and quantitative: Where tasks have a physical foundation, it is possible to derive exact relationships between morphology and performance from first principles.

**Figure 1.**
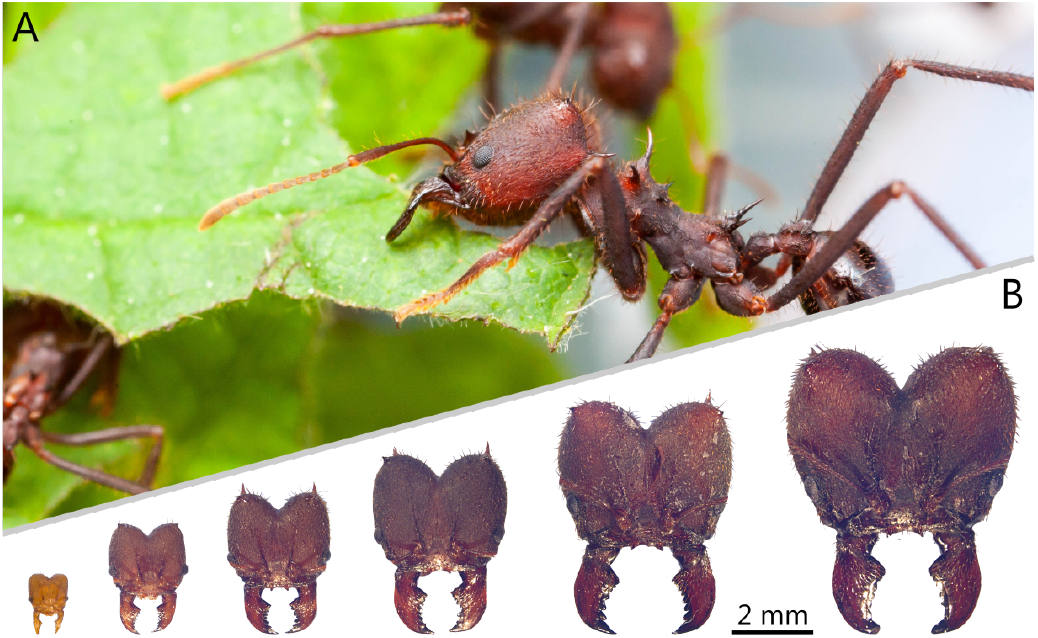
(**A**) *Atta vollenweideri* leaf-cutter ants cut and carry fresh plant fragments using their mandibular bite apparatus to supply a fungus used as crop (photo: Samuel T. Fabian). (**B**) This task is performed by workers which may vary by approximately two orders of magnitude in body mass, as illustrated here with photographs of head capsules representing this size-spectrum (dorsal view).

A key mechanical task arising in leaf-cutter ants is to cut leaf- or fruit-fragments to supply and maintain a fungus used as crop [see Fig. 1; 20–22]. Leaf-cutting occurs on an almost industrial scale, and at significant metabolic cost: Leaf-cutter ants are responsible for up to 15 % of the defoliation in the Neotropics [18, 23, 24], and the aerobic scope of leaf-cutting is comparable to that of insect flight [25]. It thus appears both plausible and necessary that leaf-cutter ants show morphological adaptations which render them particularly apt at cutting plants.

In all insects with chewing mouthparts, bite forces are generated by large muscles located in the head capsule, and transmitted to the cutting edge of the mandible via an apodeme and a mandibular joint [26–28]. Because this musculoskeletal bite system is both of behavioural, ecological and evolutionary relevance *and* can be analysed with first principles, it has received increasing attention from biomechanists [29–33], evolutionary biologists [34–39], functional morphologists [32, 40–48] and (behavioural) ecologists alike [28, 49–56]. Concretely, for an isometric contraction at zero fibre stretch, the force exerted at any point of the mandible, *F_b_*, may be written as the product between the ratio of muscle volume *V_m_* and the average fibre length *L_f_* (the physiological cross-sectional area of the muscle, *A_phys_* = *V_m_/L_f_*), the muscle stress *σ_m_*, the cosine of the pennation angle *φ*, and the mechanical advantage *MA* [28, 40, 49, 51]:

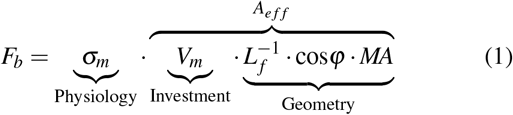

Parametrising the bite force in this way has two distinct advantages. First, it illustrates that *F_b_* may be modulated by changing one of three distinct determinants: muscle *physiology* (represented by muscle stress), total muscle *investment* (represented by muscle volume), and the *geometry* of the musculoskeletal apparatus, defined by the arrangement of muscle fibres and the lever system. The product between the latter two terms may be interpreted as the effective cross-sectional area *A_e f f_* of a muscle which acts directly at the point of force application; *A_e f f_* is thus a suitable proxy for bite force capacity.

Second, because *A_e f f_* is defined entirely by morphological quantities, it is possible to quantitatively link both size- and shape-variation across workers to a specialisation in terms of bite force capacity. Concretely, the parsimonious assumption of geometric similarity implies *V_m_* ∝ *m*, *L_f_* ∝ *m*^1/3^ and cos*φ* ∝ *MA* ∝ *m*^0^, so that *A_e f f_* ∝ *m*^2/3^ (where *m* is body mass). Any deviation from this prediction indicates shape-variation that correlates with a modulation of bite force capacity. Here, we use this predictive model as a quantitative guideline for a morphometric analysis of the bite force apparatus of *Atta vollenweideri* leafcutter ants across the entire size range. We (i) investigate if bite force capacity is modulated solely by size or also by shape differences; (ii) analyse if and why such specialisation may occur predominantly via changes in muscle investment vs. geometry; and (iii) discuss how any specialisation requires a variation of the external and internal head morphology, due to geometric and mechanical constraints.

## Materials & methods

### Study animals

Individual ants were sampled from a colony of *Atta vollenweideri*, founded and collected in Uruguay in 2014. The colony was kept in a climate chamber at 25 °*C*, 60 % relative humidity, in a 12/12 h light and dark cycle (FitoClima 12.000 PH, Aralab, Rio de Mouro, Portugal), and was fed with bramble and honey water *ad libitum*. About 30 ants, across the size spectrum, were collected from the foraging area and weighed (Explorer Analytical EX124, max. 120 g × 0.1 mg, OHAUS Corp., Parsippany, NJ, USA). From these 30 ants, we then selected 16 individuals including the heaviest (43.3 mg) and lightest (0.3 mg) specimen to achieve an approximately even spacing between log-transformed masses. The sample size was limited by the time-intense segmentation process. However, the consistently high *R*^2^ values of our scaling regressions, and the narrow confidence intervals of the associated slopes suggest that our conclusions are robust (see results).

Ants were sacrificed by freezing and decapitated using a razor blade. In order to facilitate fixative penetration, the antennae and labrum were removed with fine forceps (Dumont #5, Montignez, Switzerland), and about five holes were pierced into the head capsule using ‘00’ insect pins for specimens >1 mg. All heads were fixed in a paraformaldehyde solution (4 % in PBS, Thermo Fisher Scientific, Waltham, MA, US) for 18 h and subsequently stored in 100 % ethanol.

### Micro computed-tomography and tissue segmentation

Computed-tomography (CT) scans were performed at the Imaging Cluster at the KIT light source (see SI for details). Prior to segmentation, the tomographic image stacks were preprocessed using ‘Fiji’ [57]. Each stack was aligned such that the lateral, dorso-ventral and antero-posterior axes coincided with the principal axes of Fiji’s coordinate system (see Fig. 2). The lateral and dorso-ventral axes were defined based on the bilateral symmetry of the heads. The antero-posterior axis was defined as the line connecting the distal end of the dorsal mandible with the centre of the posterior head opening in lateral view. The oriented image stacks were imported in ‘ITK-SNAP’ [v3.6, 58] for threshold-based segmentation. Head capsule, mandibles, as well as mandible opener and closer apodemes and muscles were separated with the semi-automatic ‘active contour’ mode; directly- and filament-attached muscles were segmented as individual tissues (see below). The grayscale thresholds, set individually for each tissue, were kept constant within a sample, but were manually adjusted between samples to account for variation in contrast and brightness. For each individual, a second segmentation was performed to quantify characteristic linear dimensions of the head capsule and its volume. Artificial segmentation boundaries were added where the head capsule was open: At the anterior side, the segmentation was ‘cut-off’ at the mandible joints (see below); the posterior head opening was closed at the lateral edge of the postocciput [see 47].

**Figure 2.**
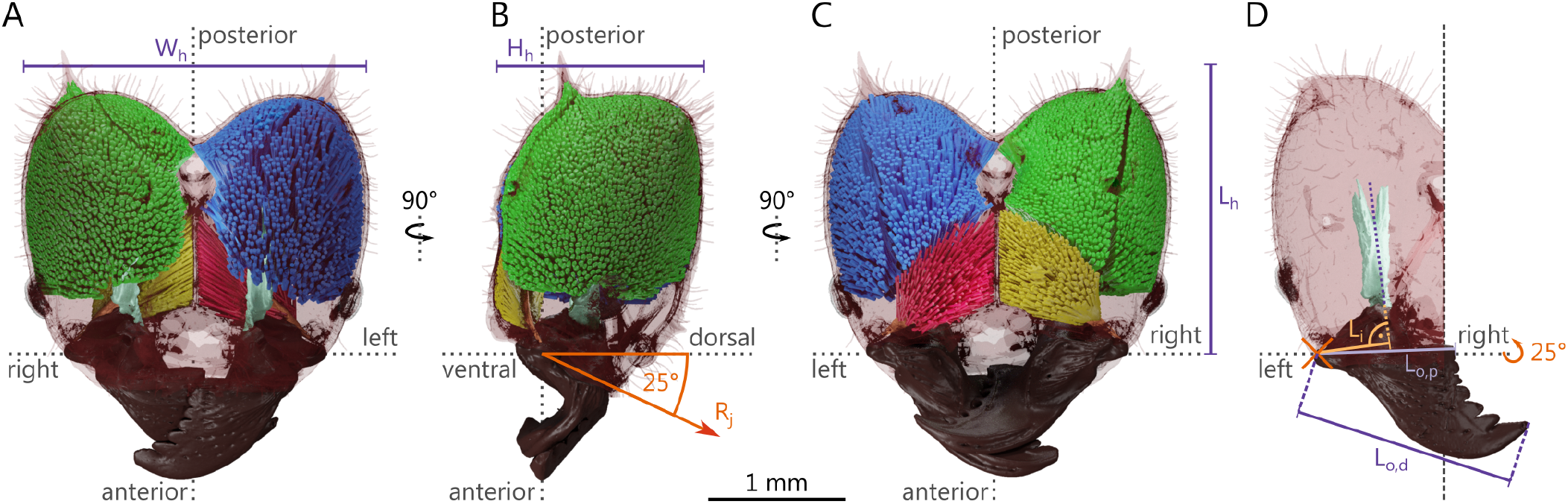
(**A**) Dorsal, (**B**) lateral and (**C**) ventral view of the internal head anatomy of a medium-sized *Atta vollenweideri* worker (10.9 mg). Tissues as segmented from micro-computed tomography scans are shown in green (closer muscle), yellow (opener muscle), turquoise (closer apodeme) and light brown (opener apodeme). Muscle fibres as reconstructed with a custom fibre-tracking algorithm are coloured in blue (closer muscle) and red (opener muscle); dashed lines indicate the lateral (left-right), antero-posterior and dorso-ventral axes. We also indicate head width *W_h_*, height, *H_h_* and length *L_h_*. (**D**) The effective inlever *L_i_*, proximal and distal outlevers, *L_o,p_* and *L_o,d_*, respectively, are defined as the perpendicular distances to the rotational axis of the mandible joint *R_j_*.

### Morphometric analysis

We extracted a series of parameters from the CT scans in order to characterise changes in the key morphological determinants of bite force, and associated changes in head and apodeme morphology.

#### Head dimensions

Head width and height were extracted from the ‘head volume’ segmentation, using Fiji’s native 2D particle analysis on all slices perpendicular to the antero-posterior axis. Based on the bilateral symmetry of the heads, head width and height were defined as the width or height of the bounding box with maximum lateral or dorso-ventral expansion, respectively. Head length, in turn, was defined as the antero-posterior distance between the first and last image slice containing segmented head volume.

#### Volume occupancy

Segmentation volumes were directly exported from ‘ITK-SNAP’. We measured the ‘functional volume’ occupied by muscle as the volume of a convex hull surrounding the segmented muscle tissue, extracted with the ‘3D convex hull’ plugin in Fiji [59]. The functional volume encapsulates all muscle fibres and the space between them including filaments and apodeme. Volume occupancy was defined as the ratio between functional volume and head volume.

#### Apodeme dimensions and orientation

The main axis of the apodeme was obtained via principle component analysis on the *xyz* coordinates of the centres-of-area of the apodeme cross-sections perpendicular to the antero-posterior axis. The apodeme image stack was then rotated such that it was perpendicular to the apodeme main axis using ‘TransformJ’ [60]. Apodeme length was defined as the distance between first and last image slice containing segmented apodeme; cross-sectional areas and perimeters were extracted via 2D particle analysis for each slice. Apodeme centre-of-mass and surface area were extracted with Fiji’s native 3D particle analysis routines.

#### Muscle fibre dimensions and orientation

Length and orientation of segmented muscle fibres were extracted with a custom tracking algorithm written in Python 3.7.6 [61]. A detailed description of the algorithm is provided in the SI. In short, the origins of individual muscle fibres were identified by dilating the head capsule, and extracting the intersections with the muscle tissue. These intersections were used as ‘seed points’ to grow fibres in an iterative process, designed to maximise fibre length and homogeneity. We extracted muscle fibre length and pennation angle for a total of 62,380 fibres across all samples (see SI for details).

Fibre pennation angle was defined as the angle between the fibre and the main axis of the apodeme [51]. Implicit in this definition is the assumption that muscle contraction moves the apodeme along its main axis. This assumption is supported by the fact that the main axis approximately coincides with the resulting muscle force vector, which has been used as an alternative reference axis for the calculation of pennation angles in previous studies [29, 32]. Pennation angles, averaged for each muscle tissue, were about two degrees smaller using this definition, but this difference was independent of worker size [Analysis of covariance (ANCOVA): F_1,56_ = 0.10, P = 0.75], within the accuracy of the angle estimation (see below), and thus does not affect conclusions on the scaling of the average pennation angle.

The muscle physiological cross-sectional area *A_phys_* was defined as the ratio between segmentation volume and average muscle fibre length. Note that all morphological muscle parameters are from scans with closed mandibles and maximum muscle contraction. Thus, closer muscle fibre are shorter, have higher pennation angles, and larger physiological crosssectional areas than in the uncontracted state [see 51].

#### Mandible dimensions

Mandible inlever was defined as the distance between (i) the insertion of the apodeme ligament to the mandible and (ii) the mandible joint (see Fig. 2D). The apodeme ligament is the segment between the anterior cranio-mandibular muscle apodeme (henceforth referred to as closer apodeme) and the mandible base [see 62, 63]; the location of the mandible joint was defined as the centre of the ridge between the ventral mandibular articulation and the atalar acetabulum [see 47]. In contrast to the inlever, the outlever is not solely defined by anatomy, but can be adjusted to serve behavioural needs. For example, leaf-cutter ants may use only the distal end of their mandibles to anchor or cut through the leaf lamina, or engage the entire blade when cutting thicker veins [19, 64]. The minimum and maximum functional outlevers were defined as the distance between point (ii) and the tip of the most proximal or distal mandibular tooth, respectively.

We then estimated the effective in- and outlevers, *L_i_*, and *L_o_*, respectively, defined as the moment arms perpendicular to the axis of rotation (and the apodeme main axis for the inlever, see Fig. 2D). This calculation required two assumptions on the kinematics of the mandibles: (i) the mandible joint is a revolute joint [44, 46, 47, 52, 65–67], and (ii), the joint rotational axis has a constant relative orientation with respect to a head-fixed coordinate system [see 68]. The second assumption was supported by preliminary kinematics analysis suggesting that the rotational axis is approximately parallel to the sagittal plane, and forms an angle of about 25 ° to the dorso-ventral axis (see Fig. 2). In comparable studies on cockroaches, dragonflies and stag beetles, the axis of rotation was deduced from morphology alone [29, 31, 32, 36, 37]. Ants, however, posses more complex joints, rendering morphological inference difficult [but see 46–48]. We observed no obvious changes in joint morphology across sizes, so that any errors introduced by this estimation are unlikely to result in a systematic effect on our scaling analysis. The mechanical advantage *MA* was defined as the ratio between effective in- and outlevers.

### Error analysis

The accuracy of the fibre identification and tracking procedure is defined by four distinct sources of error: (i) identification error, (ii) intrinsic error, (iii) segmentation error, and (iv) tracking error. We assessed these errors by comparing the result of the tracking algorithm with manual measurements of length and orientation of a subset of 278 fibres across the ant size range (see SI for details).

#### Identification error

In order to track fibres, they need to be correctly identified. Our approach based on head capsule dilation may identify too many fibres – by considering noise as a seed point – or too few, by assigning the origin of multiple fibres to one single seed point. As it was unfeasible to manually count fibres, we quantified this error indirectly, by comparing the average physiological cross-sectional areas per fibre, *A_f_*, calculated in two different ways. First, we rotated the image stacks of manually tracked fibres to be perpendicular to the fibre’s long axis (see above and SI); we then measured the cross-sectional area of these fibres by placing an ellipse around the fibre at three locations along its length. Second, we estimated *A_f_* as the ratio between the total physiological cross-sectional area and the estimated numbers of fibres (this definition conflates identification and length estimation errors, see below). An ANCOVA on log-log transformed data showed neither a significant difference between the two methods, nor an effect of size [method: F_1,38_ = 1.11, P = 0.30, interaction: F_1,38_ = 3.82, P = 0.06]. Thus, identification error is negligible.

#### Intrinsic error

Even a perfectly tracked fibre is limited in accuracy by an error intrinsic to the tracking method: the fibre with the highest quality connects the *centre* of the fibre origin at the head capsule with the *edge* of the fibre end - the longest line that contains only segmented tissue. As a result, both the fibre orientation and the fibre length have an accuracy which depends on the aspect ratio, *ρ*, of the fibres, defined as the fibre radius divided by fibre length. The fibre orientation will deviate from its main axis by *E_α_* = arctan(*ρ*), whereas the fibre length will be overestimated by a factor 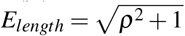. For typical muscle fibres, *ρ* ≈ 0.04, so that the angles have an accuracy of approximately 2 °, and the length is overestimated by less than one per mille.

#### Segmentation error

Because both angle and length are estimated from the segmented tissue, the accuracy is defined with respect to it. In order to quantify segmentation error, fibre orientation and length were measured manually from both segmented and unsegmented image stacks. On average, segmented fibres were 4 ± 6 % shorter than those measured directly from the CT-scans (One-sample t-test: P < 0.05). This error decreased significantly with size, approached zero for the largest workers [ANCOVA: F_1,10_ = 10.72, P < 0.01; OLS slope on semi-log data: −0.04, 95 % CI:(−0.10 | 0.01)], and is small compared to the overall variation in fibre length (about sixfold). Pennation angles did not differ significantly between unsegmented and segmented fibres (One-sample t-test: P = 0.33).

#### Tracking error

We quantified tracking error by direct comparison of manual and automated measurements. Fibres were matched between manual and automatic measurements by minimising the distance between respective fibre origins. Our custom-written algorithm overestimated the fibre length by 12 ± 21 % independent of size [ANCOVA: F_1,10_ = 1.23, P = 0.29]. Average pennation angles were overestimated by 5 ± 3 ° independent of size [ANCOVA: F_1,10_ = 0.46, P = 0.51]. These results compare favourably to the performance of commercial alternatives [69].

### Data analysis

All data analysis was performed in jupyter lab v2.1.2 [70], using R v3.6.1 [71] and Python v3.7.6 [61]. Unless stated otherwise, all values are given as mean ± standard deviation. There was no significant difference between tissues in the left and right head hemispheres [ANCOVA: P > 0.05], so we used the average. Analyses involving morphological traits, their derived quantities and body mass were performed on log_10_-transformed data. We used ordinary least squares (OLS) and standardised major axis (SMA) regressions models implemented in the R package ‘SMATR’ to describe scaling relationships [72]. The suitability of these models is subject to debate, as they involve assumptions on ‘observational’ and ‘biological’ errors [73–75]. Fortunately, the main conclusions of this paper are supported by either model. The text reports the results for OLS regressions for simplicity; SMA regression results are provided in the SI. Allometric scaling in polymorphic ants has sometimes been described with bi- or curvilinear models, i. e. a single log-log slope may not allow an adequate description of the observed allometry [6, 8, 9, 13, 76–78]. Visual inspection of key morphological traits in our data suggested an approximately linear relationship on a log-log-scale, i. e. a constant differential growth factor [79]. However, the smallest workers (head width < 1 mm), appeared to depart from this linear relationship: ‘Minims’ systematically ‘underperformed’, i. e. they had significant negative residuals (see e. g. Fig. 3D). Indeed, the assumptions of a linear model were only met when minims were excluded [assessed via a global procedure following 80], which also increased the coefficient of determination *R*^2^ considerably for all regressions on opener and closer muscle volumes, inlever and outlevers. On the basis of this statistical observation, as well as the biological argument that minims preferentially engage in brood care and gardening, and only rarely partake in foraging activities which involve cutting [7], we excluded them from all subsequent analyses.

**Figure 3.**
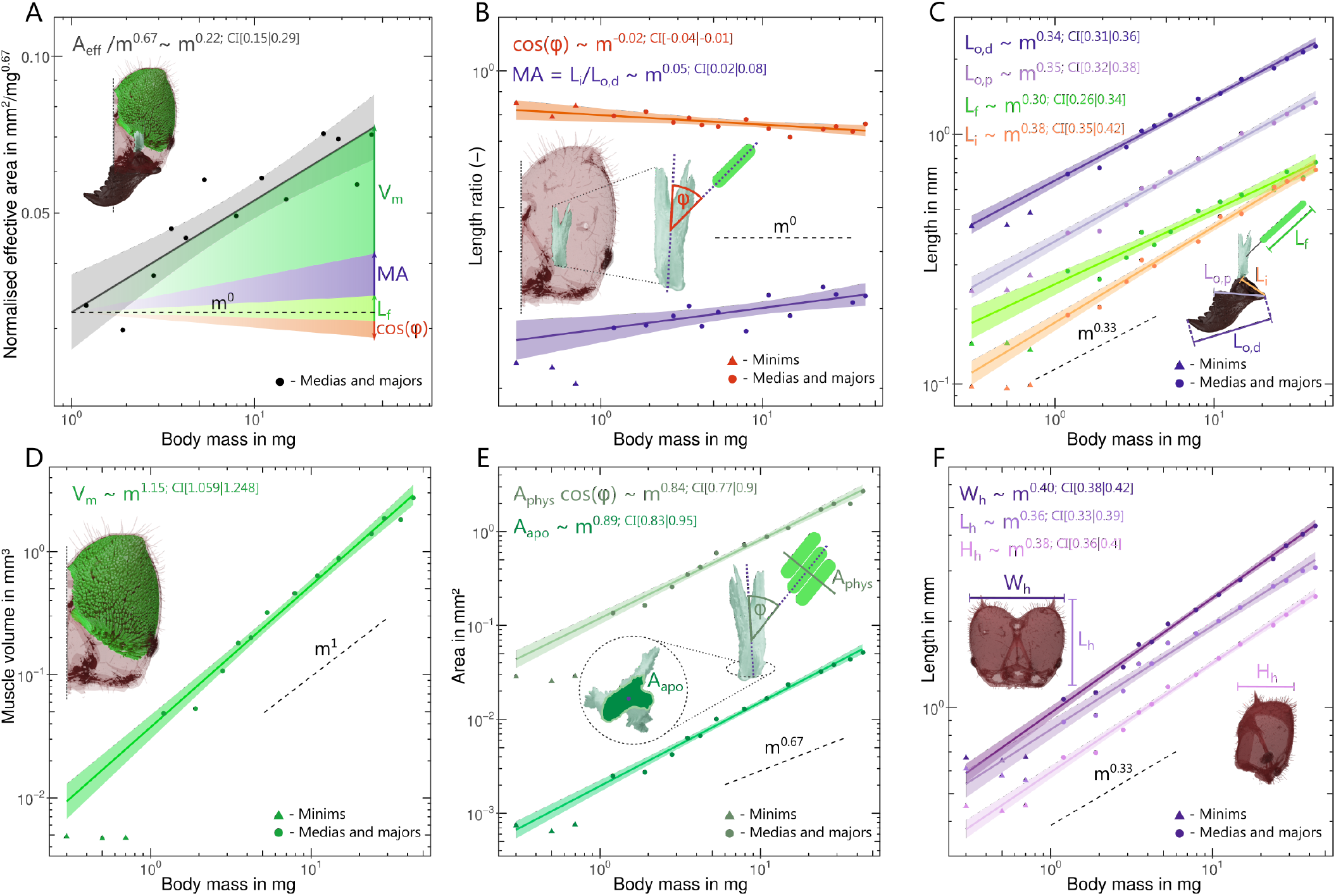
Scaling of bite force capacity, its morphological determinants and associated allometric changes in internal and external head anatomy in *A. vollenweideri* leaf-cutter ants. Solid lines show the results of ordinary least squares regressions on log-log-transformed data excluding the minims (triangles, see methods), shadings show the 95% confidence intervals, and dashed lines show predictions from isometry. (**A**) Bite force capacity is determined by investment in muscle volume *V_m_* and the geometry of the musculoskeletal apparatus, and shows strong positive allometry. The shaded areas indicate the extent to which the size-specific increase in bite force capacity is determined by the different morphological determinants. (**B**) The mechanical advantage, *MA* and cosine of the average pennation angle, cos*ϕ*, both change systematically but moderately with size. (**C**) The change in mechanical advantage is solely achieved by a positive allometry of the inlever *L_i_*; both the distal and proximal effective outlevers, *L_o,p_* and *L_o,d_*, respectively, are isometric. Fibre length, *L_f_*, shows a weak but non-significant trend to negative allometry. (**D**) Muscle volume grows with strong positive allometry, and because fibre length is isometric, this increase largely reflects an increase in the physiological cross-sectional area of the muscle. (**E**) Countering the increase in size-specific bite force associated with the positive allometry of *A_phys_*, the apodeme cross-sectional area *A_apo_* also scales with strong positive allometry, so maintaining approximately equal apodeme stress. (**F**) The increase in size-specific muscle volume is accommodated mainly by a positive allometry of head width *W_h_* and height *H_h_*; head length *L_h_* is isometric.

## Results and discussion

Cutting plant fragments is a central part of the behavioural repertoire of leaf-cutter ants. Bite force capacity is thus a biologically relevant performance metric for the suitability of an individual worker to partake in foraging. All morphological determinants of bite force capacity apart from fibre length differ significantly from isometry (see Table 1 for detailed statistics). As a cumulative result of these changes, the effective cross-sectional area, *A_e f f_* scales as *A_e f f_* ∝ *m*^0.88^ [95 % CI: (0.81 | 0.95)], in significant excess of the isometric prediction, *A_e f f_* ∝ *m*^0.67^. This difference in scaling coefficients may appear small, but it results in a substantial enhancement of the absolute bite force capacity: the largest workers have an effective cross-sectional area twice as large as predicted from changes in size alone, (45/1)^0.88−0.67^ ≈ 2; achieving this increase in bite force capacity through changes in size alone would require workers of a maximum size approximately 2^3/2^ ≈ 3 times larger than the largest workers in the colony. Two key questions emerge from this result. First, which of the ‘morphological dials’ are turned to achieve the departure from isometry? Second, what additional adaptations of internal and external head are required to implement those changes?

**Table 1.** Results of ordinary least squares regressions describing the relationship of log10-transformed determinants of bite force capacity *A_e f f_*, internal and external head dimensions with log10(body mass in mg). 95 % confidence intervals are provided in parentheses.

| Quantity | Unit | Elevation | Slope | $R^2$ |
| --- | --- | --- | --- | --- |
| $A_{eff}$ | $\text{mm}^2$ | -1.49 (-1.56, -1.42) | 0.88 (0.81, 0.95) | 0.99 |
| $V_m$ | $\text{mm}^3$ | -1.13 (-1.22, -1.03) | 1.15 (1.06, 1.25) | 0.98 |
| $L_f$ | $\mu\text{m}$ | 2.4 (2.36, 2.44) | 0.3 (0.26, 0.34) | 0.96 |
| $A_{phys}$ | $\text{mm}^2$ | -0.83 (-0.9, -0.76) | 0.86 (0.79, 0.92) | 0.99 |
| $\cos\varphi$ | (-) | -0.1 (-0.11, -0.08) | -0.02 (-0.04, -0.01) | 0.49 |
| $MA$ | (-) | -0.57 (-0.6, -0.54) | 0.05 (0.02, 0.08) | 0.54 |
| $L_i$ | mm | -0.75 (-0.79, -0.72) | 0.38 (0.35, 0.42) | 0.98 |
| $L_{o,p}$ | mm | -0.43 (-0.46, -0.4) | 0.35 (0.32, 0.38) | 0.99 |
| $L_{o,d}$ | mm | -0.19 (-0.21, -0.16) | 0.34 (0.31, 0.36) | 0.99 |
| $V_h$ | $\text{mm}^3$ | -0.56 (-0.63, -0.49) | 1.15 (1.08, 1.21) | 0.99 |
| $W_h$ | mm | -0.02 (-0.04, 0) | 0.4 (0.38, 0.42) | 0.99 |
| $H_h$ | mm | -0.23 (-0.25, -0.21) | 0.38 (0.36, 0.4) | 0.99 |
| $L_h$ | mm | -0.08 (-0.11, -0.04) | 0.36 (0.33, 0.39) | 0.98 |
| $A_{apo}$ | $\text{mm}^2$ | -2.71 (-2.77, -2.65) | 0.89 (0.83, 0.95) | 0.99 |

**Table 2.** Results of standardised major axis regressions describing the relationship of log_10_-transformed determinants of bite force capacity *A_e f f_*, internal and external head dimensions with log_10_(body mass in mg). 95 % confidence intervals are provided in parentheses.

| Quantity | Unit | Elevation | Slope | R<sup>2</sup> |
| --- | --- | --- | --- | --- |
| $A_{eff}$ | mm<sup>2</sup> | -1.5 (-1.57, -1.43) | 0.89 (0.82, 0.96) | 0.99 |
| $V_m$ | mm<sup>3</sup> | -1.13 (-1.23, -1.04) | 1.16 (1.07, 1.26) | 0.98 |
| $L_f$ | μm | 2.39 (2.35, 2.44) | 0.3 (0.26, 0.35) | 0.96 |
| $A_{phys}$ | mm<sup>2</sup> | -0.83 (-0.9, -0.76) | 0.86 (0.8, 0.93) | 0.99 |
| $\cos\phi$ | (-) | -0.09 (-0.1, -0.07) | -0.03 (-0.05, -0.02) | 0.49 |
| $MA$ | (-) | -0.58 (-0.61, -0.55) | 0.06 (0.04, 0.1) | 0.54 |
| $L_i$ | mm | -0.76 (-0.79, -0.72) | 0.39 (0.36, 0.42) | 0.98 |
| $L_{o,p}$ | mm | -0.43 (-0.46, -0.4) | 0.36 (0.33, 0.38) | 0.99 |
| $L_{o,d}$ | mm | -0.19 (-0.21, -0.16) | 0.34 (0.32, 0.36) | 0.99 |
| $V_h$ | mm<sup>3</sup> | -0.57 (-0.63, -0.5) | 1.15 (1.09, 1.22) | 0.99 |
| $W_h$ | mm | -0.02 (-0.04, 0) | 0.4 (0.38, 0.43) | 0.99 |
| $H_h$ | mm | -0.23 (-0.25, -0.21) | 0.38 (0.36, 0.4) | 0.99 |
| $L_h$ | mm | -0.08 (-0.11, -0.05) | 0.37 (0.34, 0.4) | 0.98 |
| $A_{apo}$ | mm<sup>2</sup> | -2.71 (-2.78, -2.65) | 0.9 (0.84, 0.96) | 0.99 |

### Functional constraints on mechanical advantage and pennation angle favour positive allometry of *A_phys_*

The maximum bite force capacity is determined by the investment in muscle volume *V_m_*, and by the average fibre length *L_f_*, pennation angle *φ* and the mechanical advantage *MA* of the force transmission system (see Eq. 1). About 70 % of the observed disproportionate increase in *A_e f f_* is achieved by a disproportionate increase in muscle volume *V_m_*. *MA* and *L_f_*, in turn, make a contribution of about 40 %; the systematic decrease of *φ* seemingly reduces size-specific bite force capacity by about 10 % (see Table 1 and Fig. 3A). Thus, bite force capacity is substantially enhanced through an increased investment in muscle volume as opposed to adjustments in the geometry of the musculoskeletal apparatus. What are the advantages and disadvantages of altering the relative force capacity through a systematic increase in volume investment over changes in geometry, and what constraints may limit one strategy or favour another?

#### Mechanical advantage

Across worker sizes, *MA* shows limited but significant positive allometry; *MA* increases by about 20%, irrespective of whether it is defined with respect to the distal or proximal sections of the gnathal edge (*sensu* [81]; see Table 1 and Fig. 3B). A systematic change in *MA* may be achieved by an increase of the effective inlever (*L_i_*), a decrease in the effective outlever (*L_o_*), or a combination of both. In *A. vollenweideri*, the positive allometry in *MA* is solely driven by the first option (see Fig. 3C). Although halving *L_o_* would result in the same numerical change of *MA* as doubling *L_i_*, these two changes are not functionally equivalent: An increase in *L_i_* increases the force available at *any* point along the mandibular cutting edge. In contrast, a shortening of *L_o_* merely reduces the functional length of the mandible without providing a clear functional advantage – smaller outlevers can also be achieved by simply biting with a more proximal part of the mandible. Because this behavioural flexibility is not afforded to *L_i_* – which is anatomically fixed it is functionally sensible to drive systematic changes in *MA* through changes in inlever length.

Notably, both the absolute value of the *MA* and its variation, 0.27 < *MA* < 0.32 for the distal outlever, are at the lower end of values reported across numerous insect taxa [typically < *MA* < 0.8, see 82]. Thus, the ants appear to utilise only a small fraction of the theoretically available scaling capacity a change from 0.3 to 0.8 across worker sizes would result in *MA* ∝ *m*^0.26^, a factor of five in excess of the observed scaling. We argue that the scaling capacity of *MA* is constrained for at least two reasons. First, systematic increases in *L_i_* are difficult to implement, because inlevers are internal to the head-capsule. Second, the magnitude of the bite force is not the sole functional determinant of bite performance. Instead, ant workers may have to maintain an approximately similar opening range across sizes, which results in a functional coupling of in- and outlever. To retain an equivalent mandibular opening range, a strong positive allometry of *MA* would require relatively longer muscle fibres, or an increase in characteristic muscle strain. Implementing either option likely requires substantial changes to head anatomy or muscle physiology, and may therefore only be possible across more distantly related species, where developmental and phylogenetic constraints are relaxed. Indeed, the *MA* of the mandibular system contains significant phylogenetic signal [82], and extreme variations in outlever such as between male and female stag beetles are matched by corresponding changes in inlever [29], both consistent with this interpretation.

#### Pennation angle

Across worker sizes, the cosine of the average pennation angle *φ* decreases significantly by about 8%, i. e. *φ* increases from about 37 ° to 43 ° (see Table 1 and Fig. 3B). Thus, the systematic change in *φ* seemingly *decreases* the bite force capacity of larger workers. However, this conclusion is premature. Pennate muscle fibres require an attachment area which exceeds their cross-sectional area by a factor *csc*(*φ*) [49, 51, 52, 83–85]. This dependency of the physiological cross-sectional area on *φ* introduces a sin-term into Eq. 1, so that the maximum bite force capacity occurs not for *φ* = 0°, as may be concluded from Eq. 1. Instead, the maximum occurs for 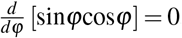, i. e. for *φ* = 45°, remarkably close to the values measured for the largest workers [also see 51]. Concretely, although the fraction of the force acting along the apodeme decreases as *m*^−0.02^, the potential for an increase in *A_phys_* scales as *m*^0.03^ so that the net result of the increase in *φ* may well be an increase in bite force capacity; we discuss how the disproportionate increase in *A_phys_* is implemented below. Because small workers already have average pennation angles close to the theoretical optimum for force capacity, the net change in bite force capacity arising from changes in *φ* is negligible. Indeed, the similarity of *φ* across sizes likely reflects a f unctional limitation in analogy to the constraint imposed on *L_i_*: changes in *φ* at constant size-specific muscle length would either alter the functional opening range of the mandible, or require a systematic variation in characteristic muscle strain [see 84, 86].

#### Muscle length and volume

The segmented volume of the closer muscles fibres scales with positive allometry, *V_m_* ∝ *m*^1.15^. In contrast, the length of the muscle fibres scales close to isometry, *L_f_* ∝ *m*^0.30^ (see Table 1 for detailed statistics). Thus, the positive allometry of *V_m_* exclusively reflects a strong positive allometry of *A_phys_* ∝ *m*^0.86^, and hence bite force capacity. Leaf-cutter ants thus deploy a ‘hybrid strategy’ of combining positive allometry of muscle volume with isometry of fibre length, simultaneously satisfying two biological demands: the differential growth of *L_f_* and 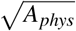 systematically increases bite force capacity, and the isometric growth of *L_f_* ensures that mandibular opening range remains approximately size-invariant.

### Positive allometry of bite force capacity requires substantial changes to external and internal head anatomy

We have demonstrated that larger *A. vollenweideri* leaf-cutter ants boost their bite force capacity via an increase investment in muscle volume, while the geometric arrangement defined by muscle length, pennation angle and the mechanical advantage only shows minor size-specific changes. This positive allometric growth results in a substantial increase of bite force capacity (see Fig. 3A), but poses a significant challenge for external and internal head anatomy: Internal adaptations are required to provide the attachment area for a relatively larger *A_phys_*; external adaptations are necessary to combine the positive allometry of *A_phys_* with isometry of *L_f_*, and to provide sufficient space for other functional tissues.

#### Internal anatomy

The disproportionate increase in *A_phys_* needs to be matched by an equivalent increase of the internal attachment area. In order to model this increase, we approximate the shape of each half of the head capsule as a cylinder with radius *R* and height *h* = *f_h_R* (with *f_h_* ≥ 0). This cylinder is terminated with a spherical cap, also with radius *R*, at its posterior end. The internal attachment area, in turn, is defined by a cylindrical part with equal height *h*, but radius *r* = *f_r_R* for the cylindrical section and the spherical cap (with 0 ≤ *f_r_* ≤ 1. The internal area is not equal to the apodeme surface area, because most muscle fibres attach via filaments, see Fig. 4A). We estimate *h* as the apodeme length, *R* as a quarter of the head width, and *r* = *R − L_f_*sin*φ*. This simple model predicts the functional volume occupied by muscle to 3 % accuracy (*V_f unc_* ≈ *V_ext_*), and *A_phys_* to 19 % accuracy (*A_phys_* ≈ sin*φA_int_*). Having demonstrated that our geometric approximation captures the salient features of the internal and external head geometry pertaining to the arrangement of the closer muscle, we turn our attention to two functional predictions it enables.

**Figure 4.**
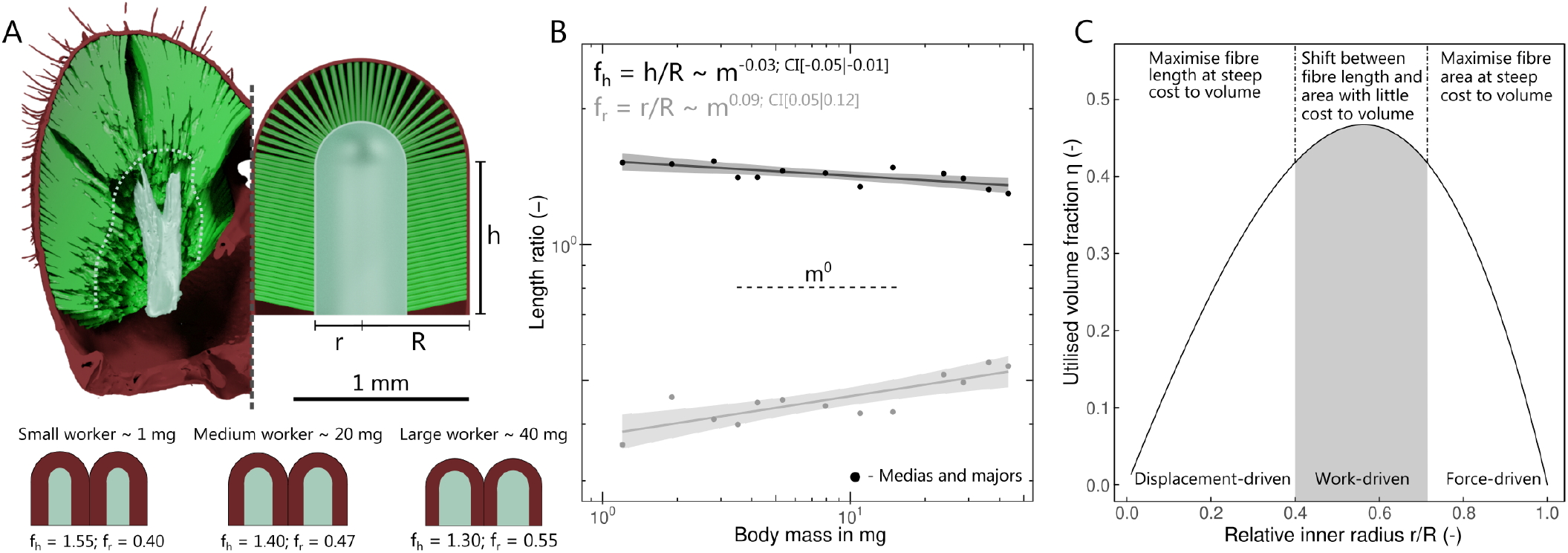
(**A**) The volume of half a head capsule may be modelled as a cylinder of length *h* and radius *R*, terminated by a spherical cap. The internal fibre attachment area surrounds and internal volume of identical geometry, but with radius *r* ≤ *R*; this volume contains both the apodeme, and muscle-filaments (see main text). (**B**) The ratio *h/R* is negatively allometric so that heads of larger ants appear more ‘heart-shaped’. The ratio *r/R*, in turn, is positively allometric, indicating a change in the surface to volume ratio of the attaching muscle (see main text). (**C**) The internal head morphology of *A. vollenweideri* leaf-cutter ants achieves approximately optimal volume utilisation (here shown for a mean value of *h/R* = 1.4), and lies in a morphological region where fibre length and area can be ‘exchanged’ without a significant variation in utilised volume.

First, we note that, although *A_phys_* can be increased by increasing either *h* or *r*, deviating from the isometric prediction of 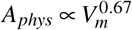 requires a systematic variation in *r/R* = *f_r_*, which controls the surface to volume ratio of the muscle (see SI):

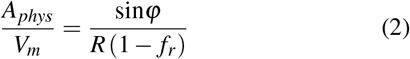

Indeed, in *A. vollenweideri*, *r* grows more quickly than *R*, so that *f_r_* increases systematically with size [*r* ∝ *m*^0.49^. 95 % CI: (0.45 | 0.52); *f_r_* ∝ *m*^0.09^. 95 % CI: (0.05 | 0.12), see Fig. 4B]. As a consequence of this shift in internal anatomy, the positive allometry of muscle volume is disproportionally invested into *A_phys_* instead of *L_f_*. Additional support for the hypothesis that a disproportionate increase in *A_phys_* is the dominant objective is provided by the observation that *h* grows more slowly with mass than *r*, [*h* ∝ *m*^0.37^. 95 % CI: (0.34 | 0.39)].

Second, we seek to demonstrate that heads of *A. vollenweideri* workers occupy a morphological space where a shift of volume from fibre length to area can be achieved without a significant variation of the total muscle volume. The origin of the complex relationship between length, area and volume is rooted in the space-constraint imposed by an exoskeleton: The area occupied by a cross-section through a muscle fibre is independent of its length, but the internal area available to attach it decreases as the fibre gets longer [51, 52]. For a given head volume, there thus exist a trade-off between maximising fibre length or fibre cross-sectional area: At one extreme are fibres of maximum length which have no internal area to attach to; at the other extreme are fibres with a maximum *A_phys_* equal to the internal surface area of the head capsule, but of miniscule length. The *volume* of the muscle, then, is maximised at some compromise between investment in fibre length vs area. In order to calculate the geometric arrangement which optimises muscle volume, we consider the fraction of the available head volume occupied by the closer muscle, *η*, which is a function of *f_r_* and *f_h_* (see SI):

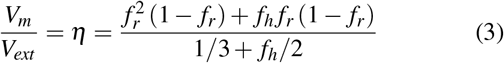

The maximum for *η* falls between two extremes set by a high-aspect ratio cylinder (*f_h_ >> f_r_*, *η_max_* = 0.5, *f_r_*(*η_max_*) = 0.5) and a hemisphere (*f_h_* = 0, *η_max_* = 0.44, *f_r_*(*η_max_*) = 0.67; see SI). Thus, the maximum volume that can be utilised is roughly half of the head capsule, which requires *f_r_* ≈ 0.5. For *Atta*-workers, our data suggest 0.36 < *f_r_* < 0.55 and 1.30 < *f_h_* < 1.53, covering a functionally meaningful range (see Fig. 4B): For *f_r_* << 0.5, muscle length is favoured over area at the steep expense of muscle volume. For *f_r_* >> 0.5, in turn, volume is sacrificed to maximise area. For *f_r_* ≈ 0.5 as observed in *A. vollenweideri*, the partitioning of volume into fibre length and cross-sectional area can be altered without a large variation in total muscle volume (see Fig. 4C): The geometric changes in internal head morphology increase the size-specific *A_phys_* by more than 50 %, but only alter *η* by about 12 %.

The internal attachment area surrounds an internal volume. In principle, this internal volume could be filled entirely with apodeme, but in insects, individual muscle fibres may alternatively attach via a single, thin, filament-like process of the apodeme [50]. The fraction of filamentvs directly attached muscle fibres differs across species, and likely reflects functional specialisation [50–52]. In *A. vollenweideri*, 98 ± 2 % of muscle fibres are filament-attached, independent of worker size [ANOVA: F_1,11_ = 1.98, P = 0.18]. What is the advantage of such a strong bias towards filament-attached fibres?

Hypotheses on the functional significance of filament-attached muscle fibres are scarce, but previous work has suggested that using filaments optimises area utilisation [51, 52]. Although filaments indeed result in an increase of the available attachment area, this is only a sufficient, but not a necessary condition: the same increase could be achieved by increasing the volume occupied by the apodeme instead. Indeed, filament-attached fibres appear to be an alternative to secondary and tertiary branching of the apodeme [51, 52], suggesting that the optimisation volume utilisation may not be the dominant driving factor for filament-attachment. We demonstrate in the SI that the use of filaments in *A. vollenweideri* substantially reduces the required cuticle volume by an amount approximately equal to the cuticle volume of the entire head capsule. Thus, filament-attachment has the advantage of significantly reducing material investment; such ‘light-weight’ construction may be particularly important in insects which operate close to maximum volume occupancy.

Filament-attachment may also alter how muscle shortening translates to apodeme motion. For example, filament-attached fibres are shorter, and thus slower [51, 52]. In addition, the relationship between apodeme displacement and pennation angle change differs between filament- and directly-attached muscle fibres of equal resting length. The resulting complex interaction between filament length, apodeme displacement, pennation angle, muscle contraction and average force is beyond the scope of this article, and will be addressed in a future study.

Filament-attachment of muscle fibres constitutes a powerful strategy to reduce material investment, but the apodeme itself cannot be arbitrarily small. Instead, apodeme size is bound by two constraints. First, the apodeme needs to have a surface area large enough to attach all muscle fibres. Only about 6 ± 4 % of the apodeme surface area is covered with either filaments or directly-attached muscle fibres (see SI). Consequently, the surface area is unlikely to be a limiting constraint for apodeme size. Second, the apodeme needs to have a cross-sectional area sufficiently large to withstand the muscle force without risking failure. This demand may be formally described as the condition that the ratio between two characteristic forces is to remain constant:

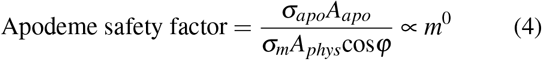

Here, *A_apo_* is the cross-sectional area of the apodeme at the location of maximum stress, and *σ_apo_* is the yield strength of the apodeme. If both characteristic stresses are size-invariant, the above condition is satisfied if the scaling of *A_apo_* is close to the scaling of *A_phys_*cos*φ* ∝ *m*^0.84^. Consistent with this condition, we find that *A_apo_* ∝ *m*^0.89^ [95 % CI: (0.83 | 0.95), Fig. 3E; see SI for how we determined *A_apo_*]. Hence, ants possess apodemes with cross-sectional areas just large enough to keep the stress approximately constant, *σ_apo_* ∝ *m*^−0.06^ [95 % CI: (−0.10 | - 0.02)]. The simple expression for the safety factor highlights a second important functional demand: the extent to which volume can be saved through the use of filament-attached fibres is modulated by the ratio *σ_apo_/σ_m_*. For representative values of *σ_m_* ≈ 0.30 *MPa*, similar to values for striated muscles in vertebrates [87], and well within the range reported for arthropods [29, 31, 88–91], and *σ_apo_* ≈ 100 − 600 MPa ([92–94] and SI), this ratio is typically *σ_apo_/σ_m_* ≈ 300 − 1800. The implemented area ratio, in turn, is cos*φA_phys_/A_apo_* = 55 ± 5, suggesting that the apodeme operates at a safety factor of ≈ 5 − 30.

#### External anatomy

The positive allometry of muscle volume poses a challenge, as larger ants need to accommodate disproportionally larger closer muscles. This challenge can be addressed by (i) increasing volume occupancy, and/or (ii), increasing the relative head volume. Volume occupancy *χ* indeed increases from 70 to over 80 % [in accordance with previous studies on ant mandible muscles, e. g. 50, 52], *χ* ∝ *m*^0.05^ [95 % CI: (0.03 | 0.06)]. This increase accounts for about 20 % of the increase in the functional volume in excess of isometry *V_f unc_* ∝ *m*^1.19^, as (45/1)^1.19−1.00^ ≈ 2. The remaining 80 % are enabled by disproportionally larger head capsules, *V_h_* ∝ *m*^1.15^ [95 % CI: (1.08| 1.21)]. Larger ants tend to have proportionally smaller brains [12, 13, 45, 77, 95], which may provide some flexibility for muscle occupancy. However, smaller ants already have the majority of their heads filled by muscle, and larger ants likely need to provide space for other tissues inside the head capsule, so that the strong allometry of *V_f unc_* must be predominantly achieved by a positive allometry of overall head volume.

The positive allometry of head volume can be modulated by scaling head length, width and height. We find that head width displays the strongest positive allometry *W_h_* ∝ *m*^0.40^ [as previously reported for *Atta*, see 7, 10, 11], followed by head height, *H_h_* ∝ *m*^0^.38. Head length, in turn, only shows a weak tendency for positive allometry *L_h_* ∝ *m*^0^.36 [95 % CI: (0.33 0.39)] (see Fig. 3F). What explains this seeming preference for increasing some head dimensions more strongly than others?

One possible explanation might lie in the expansion of the internal attachment area: changing the surface-to-volume ratio requires a strong positive allometry of the internal radius *r* ∝ *m*^0.49^ (see above). Coupled with the isometric growth of fibre length, this allometric expansion affects the relative spatial demand in width and height more strongly than it does in length, as a larger fraction of head width and height is occupied by *r*. In other words, the effect of the strong positive allometry of *r* on head length is likely attenuated by the apodeme length, and the space occupied by the opener muscle (see Fig. 2). Head width allometry, in comparison to head height, might be further driven by the need to accommodate longer mandible inlevers, which are approximately aligned with the lateral axis.

### Why vary shape to boost bite performance?

The bite force capacity in *A. vollenweideri* leaf-cutter ants shows strong positive allometry, mainly achieved by a disproportionate increase in muscle investment. Bite force capacity is of particular relevance for leaf-cutter ants, as it influences the diversity of plant material that can be processed by colony workers [7, 15, 96, 97]. Hence, our results add further support to the hypothesis that size-polymorphism in leaf-cutter ants, and the associated adaptations in shape, may enable them to forage on a broader spectrum of food sources [7, 10, 20, 98, 99]. Why alter force capacity not only via size-but also via shape-differences?

We propose three potential reasons. First, achieving the same bite force with isometry would require a worker three times heavier than the largest worker in an allometric colony workforce. The positive allometry thus increases the effective size range of the colony workforce by a factor of three, but at reduced ‘production cost’ (which is proportional to mass). It is well established that larger workers cut tougher leaves [7, 15, 96, 97, 100, 101], and our results suggest that they may even cut *relatively* tougher leaves. Second, a disproportionate increase in bite force may be required to compensate an increase in mandibular cutting force – the force required to cut a material with mandibles – which may scale in proportion to a characteristic mandible length [see 102, 103]. Third, disproportionally large heads may enable an increase in size-specific bite force capacity, but they may limit or reduce the ability of workers to perform other tasks, i. e. they render them less generalist. For example, the increased head size may reduce mobility at the nest entrance and within the nest where small heads may be beneficial to successfully manoeuvre through the close-knit structure of the fungal garden [see Fig. 3F; 104]. Indeed, significant numbers of larger workers typically only exist in colonies which exceed a minimum size [17], consistent with the hypothesis of specialised large workers but generalist small workers.

Leaf-cutter ant workers show substantial, size-specific modifications to internal and external anatomy which increase their bite force capacity. The extent to which this specialisation improves colony performance requires to integrate the morphological findings of this study with quantitative measurements of cutting performance to assess ‘efficiency’, and with behavioural assays to assess how differences in efficiency are reflected in task allocation. We hope that such work will provide a comprehensive and integrative picture of the interaction between worker polymorphism, task specialisation, and foraging efficiency.

## Acknowledgments

This study is part of a project that has received funding from the European Research Council (ERC) under the European Union’s Horizon 2020 research and innovation programme (Grant agreement No. 851705, to David Labonte). We acknowledge the KIT light source for provision of instruments at their beamlines and we would like to thank the Institute for Beam Physics and Technology (IBPT) for the operation of the storage ring, the Karlsruhe Research Accelerator (KARA). Further, we gratefully acknowledge the data storage service SDS@hd supported by the Ministry of Science, Research and the Arts Baden-Württemberg (MWK) and the German Research Foundation (DFG) through grant INST 35/1314-1 FUGG and INST 35/1503-1 FUGG.

## Supplementary Materials

### Tomography

All scans were conducted with a parallel polychromatic X-ray beam produced by a 1.5 T bending magnet. The beam was spectrally filtered by 0.7 mm aluminum. A fast indirect detector system was employed, consisting of a 12 μm LSO:Tb scintillator [105] and a diffraction limited optical microscope (Optique Peter) [106] coupled with a 12 bit pco.dimax high speed camera with 2016 × 2016 pixels. Scans were done by taking 3,000 projections at 70 fps over an angular range of 180 °. Depending on their sizes, the heads were scanned with optical magnifications of either 5× or 10×, resulting in effective pixel sizes of 2.44 μm and 1.22 μm, respectively. The control system ‘concert’ [107] was employed for automated data acquisition and online reconstruction of tomographic slices for data quality assurance during the measurement campaign. Data processing included flat field correction and phase retrieval of the projections based on the transport of intensity equation [108]. The execution of the pipelines, including tomographic reconstruction, was performed by the UFO framework [109].

### Fibre-tracking algorithm

A custom-written algorithm was used to determine the number of muscle fibres, their length and orientation. Inspired by commercial software such as ‘Amira XTracing’ or ‘Avizo XFiber’ (Thermo Fisher Scientific Inc., Waltham, Massachusetts, USA), individual fibres were grown from ‘seed points’ [see 69, 110], placed at the origin of individual muscle fibres on the head capsule. Fibre origins were identified through a series of three steps in ‘Fiji’ (see Fig. 5):

**Figure 5.**
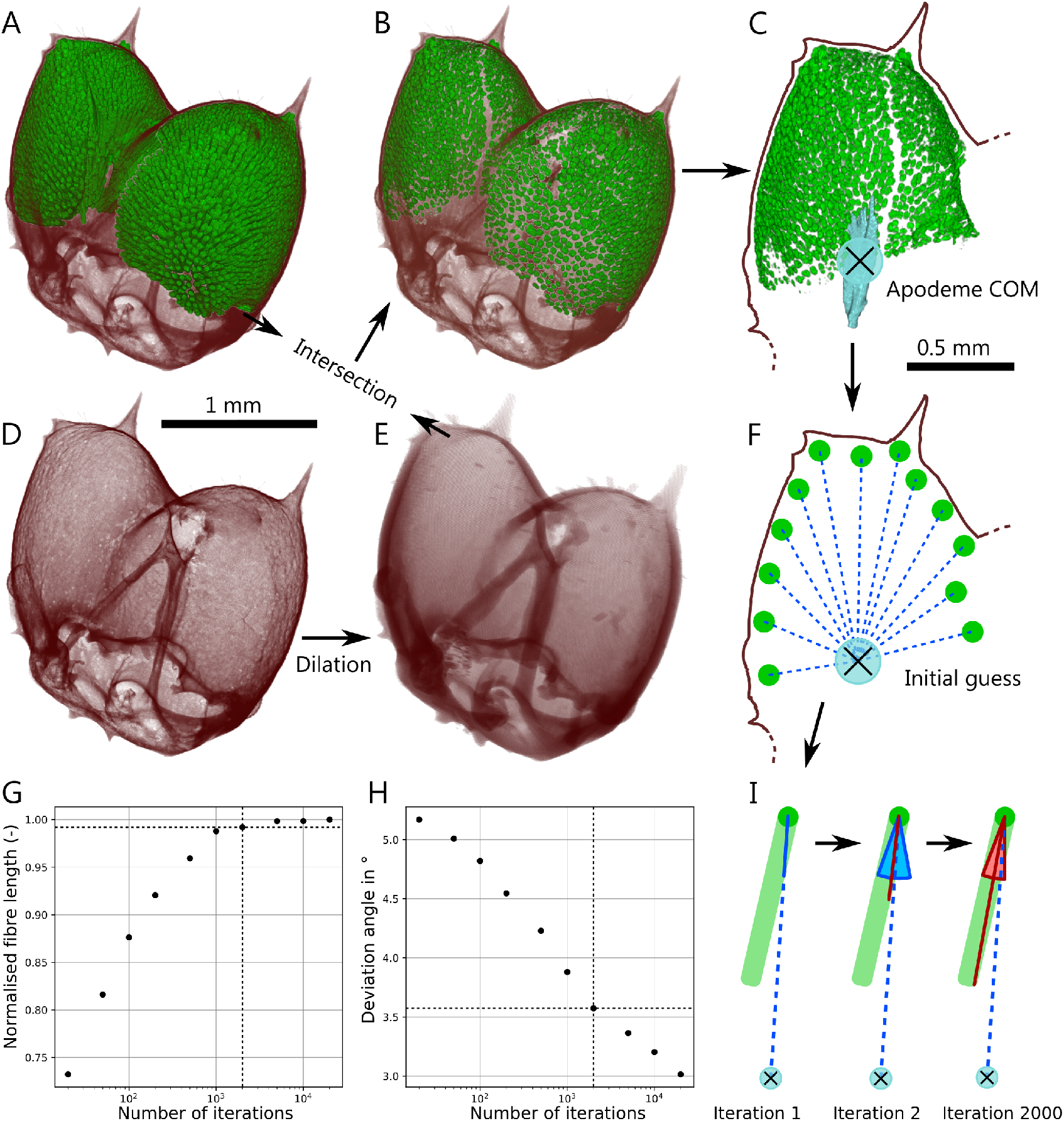
We extracted the length and pennation angle of every fibre of the closer muscle with a custom-written tracking algorithm. The head capsule segmentation (**D**) was dilated, and the intersection (**B**) between dilated head capsule (**E**) and closer muscle (**A**) was calculated, resulting in spatially distinct fibre seeds. The apodeme centre-of-mass (COM) was calculated (**C**), and the lines connecting fibre seeds and apodeme COM were selected as initial fibre orientations (**F**). This ‘initial guess’ was then refined through an iterative optimisation process (**I**), favouring long and homogenous fibre segments. Fibre length and pennation angle were extracted for a subset of fibres, revealing that fibre length increases only little (**G**), and pennation angle showed small variation for repeated measurements (**H**), after 2000 iterations of the optimisation algorithm.

First, the segmented head capsule was dilated by a factor proportional to the average muscle fibre thickness (Fig. 5 D). The number of dilation iterations *n_dil_* was defined as a constant proportion of the fibre diameter extracted from manual measurements (see below), and was interpolated across the size range using a linear regression on log-log-transformed data. Visual inspection suggested that a dilation equal to two-thirds of the diameter in pixels resulted in intersections sufficiently large to be clearly identified as individual muscle fibres, and small enough to prevent fibre merging (see Fig. 5 B).

Second, the intersections between dilated head capsule and muscle tissue were extracted (see Fig. 5 B & E).

Third, the number of distinct intersections, exceeding a minimum volume, and their centres-of-mass were calculated via a 3D particle analysis. The *xyz* coordinates of the centres-of-mass were used as the origins of these fibres (Fig. 5 F).

Starting from these seed points, we estimated both the length and orientation of the muscle fibres via an iterative ‘fibre-tracking’ process. This process takes advantage of the fact that muscle fibres in insects are approximately straight lines. Each fibre seed point was assigned an initial orientation, equivalent to the shortest line between the seed point and the centre-of-mass of the apodeme (Fig. 5 F). Each seed was then grown along this orientation pixel by pixel, until unsegmented tissue was reached. To account for irregularities in fibre morphology and segmentation, a small number of unsegmented pixels was allowed to be crossed. This number was calculated as *n_cancel_* = (*s_min_ − n_dil_*)/2, where *s_min_* represents the average minimum pixel distance between individual seed points. This cancel criterion accounts for variation in both fibre diameter and density, and usually equalled about four pixels. Occasionally, the seed point itself was unsegmented, leading to ‘zero-length’ fibres. In these cases, a new seed point was selected at random from within the bounding box of the same particle. If a fibre could be grown to a length at least two times *n_dil_*, the seed point was considered acceptable.

As the concluding step of each algorithmic iteration, the quality of the grown fibre was assessed by dividing its length with the variation in greyscale value along its length, calculated from unsegmented CT data. If the fibre was long, located within a single muscle fibre and crosses few small unsegmented gaps, the variation in greyscale was small and the quality metric large. If, in contrast, it was short, crosses numerous unsegmented gaps and includes tissue from several muscle fibres, the greyscale variation was large and the quality metric small. Hence, the algorithm favoured long and homogenous segments.

This process was conducted once for the initial orientation, and was then repeated 2000 times for orientations chosen at random from a ‘search cone’ defined with respect to the fibre which has the highest quality metric. This search cone was created by rotating the cone’s centre line around all three coordinate axes by angles taken from a population of angles normally-distributed around zero degrees. The standard deviation of this normal distribution was set to 10 ° for the first iteration and was subsequently linearly decreased with increasing number of iterations to a minimum of 0.1 ° (see Fig. 5). If the quality metric of the current iteration had increased compared to the previous ‘best guess’, the orientation of the search cone was updated. Otherwise, the next iteration was conducted along an orientation chosen at random from the previous search cone.

The number of iterations per fibre was determined by preliminary measurements using a subset of ten fibres from one closer muscle tissue. Each fibre was tracked for a number of iterations ranging between 20 and 20,000, and fibre length and pennation angle were extracted (Fibre length was measured as the number of segmented pixels along the fibre orientation multiplied with the scan resolution). This process was repeated 50 times for each fibre in order to quantify the repeatability of the outcome. After 2000 iterations the average fibre length was less than 1 % shorter compared to the maximum extracted at 20,000 iterations, and the average standard deviation of the pennation angles dropped to 3.6 °, less than twice the minimum error set by the intrinsic limit of the method (see Fig. 5). Hence, a number of 2000 iterations was regarded sufficiently large to yield reliable results while keeping the computational effort manageable.

Fibres with aspect ratios below two were removed from the dataset as such short fibres were not observed in the CT-scans, and hence likely reflect incorrectly identified seed points (see main text). Fibre length was measured as the total number of segmented pixels between the fibre seed point and point of fibre termination. This calculation does not account for a small part of the fibre between seed point and head capsule, approximately equal to half the number of dilation steps. To correct for this systematic error, another fibre was grown from the seed point in the opposite direction but along the final fibre orientation, until unsegmented tissue was reached. The length of this fibre segment was then added to the fibre length.

In order to validate the fibre-tracking algorithm, a subset of 40 opener and closer muscle fibres were manually analysed for seven ants across the entire size spectrum (278 in total – the opener muscle tissue of the smallest ant had only 18 suitable fibres). Individual fibres were identified using the results of preliminary 3D particle analysis on the muscle intersections in Fiji. Fibres were selected for tracking when distinctly visible in both CT-scan and segmentation image. The locations of the fibre centres at origin and insertion to the apodeme were extracted, and fibre length and orientation were calculated (see above and main text).

### Cuticle saving via filament-attachment

We estimated the material volume saved by filament-attachment as follows: The total volume of filament is *V_f il_* = *n _f_ A_f il_ L_f il_*, where *n _f_* is the number of filament-attached fibres, *L_f il_* the average filament length (estimated from internal head dimensions), and *A_f il_* their cross-sectional area (calculated assuming that the cross-section approximates a circle of diameter *d*). Filaments were too thin to be reliably segmented, but visual inspection of the CT scans suggested that *d* ≈ 4 *μm* with little variation within and across specimens [also see 50]. As almost all muscle fibres in *A. vollenweideri* were attached via filaments, the amount of internal cuticle required for filament-attached fibres then equals the sum of *V_f il_* and the apodeme volume *V_apo_*. In contrast, the amount of internal cuticle required for direct attachment is equal to the entire internal volume *V_int_*. The difference between these volumes, the volume saved through filament-attachment for both head halves is *V_saved_* = 2(*V_int_* − (*V_f il_ + V_apo_*)), which was approximately equal to the cuticle volume of the entire head capsule. Hence, filament-attachment substantially reduces the required cuticle tissue, and might be a light-weight strategy to satisfy the excessive demand of fibre attachment area.

### Area coverage of apodeme

Because the majority of muscle fibre attach to the apodeme via filaments, the required surface area is dominated by the relatively small cross-section of the filaments. We estimated the area fraction covered as (*n_f_ A_f il_* + *n_d_ A_f_*)/(*sin*(*φ*)*A_apo,s_*), where *A_apo,s_* is the apodeme surface area, and *n_d_* corresponds to the number of directly-attached fibres.

### Estimation of *A_apo_* and *σ_apo_*

The maximum force is experienced by the anterior parts of the apodeme cross-section. Accordingly, the cross-sectional area of the apodeme is largest close to mandible insertion and decreases towards its posterior end [see 52]. In order to identify the location of maximum stress, an assumption on the increase of force with apodeme length is required. The simplest plausible assumption is that the number of fibres increases along the length of the apodeme in proportion to the apodeme perimeter. The predicted increase in force is larger than the increase in cross-sectional area, so that the highest stresses is expected at the anterior end of the apodeme, where the cross-sectional area is maximal. Maximum cross-sectional area scales with strong positive allometry *A_apo_* ∝ *m*^0.89^ [95 % CI: (0.83 | 0.95), see Fig. 3E], similar to the increase of maximum load along the apodeme, which scales proportional to *A_phys_cos*(*φ*) ∝ *m*^0.84^. A similar conclusion holds if the average cross-sectional area is used instead, 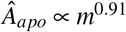 [95 % CI: (0.85 | 0.97).

In order to estimate the approximate tensile strength of the apodeme, we note that its density appears similar to that of the head capsule, which has a modulus of the order of 6 GPa (F. Puffel, unpublished data). The strength may then be estimated as approximately 1/60 of the modulus [94], i. e. *σ_apo_* = 100 MPa. Alternatively, we may consider available estimates on the longitudinal tensile strength of insect cuticle which is approximately 94 MPa [92], or the tensile strength of the leg extensor apodeme in locusts, which is 600 MPa [93]. These estimates change the conclusion on the numerical value of the safety factor, but do not affect the conclusion that the apodeme strength is at least a hundred times larger than the muscle stress.

### A simple geometric model of the ant head capsule

We model the shape of each half of the head capsule as a cylinder with radius *R* and height *h* = *f_h_R* (with *f_h_* ≥ 0). This cylinder is terminated with a spherical cap, also with radius *R*, at its posterior end. The internal attachment area, in turn, is defined by a cylindrical part with equal height *h*, but radius *r* = *f_r_R* for the cylindrical section and the spherical cap (with 0 ≥ *f_r_* ≥ 1. The external volume is *V_ext_* = *πR*^2^*h* + 2/3*πR*^3^, the internal attachment area is *A_int_* = 2*πr* (*h* + *r*), and the volume available for muscle fibres is *V_m_* = *A_int_* (*R − r*). It is crucial to note that this volume is not equal to the difference between external and internal volume, because the internal and external areas are not equal. *A_phys_* can be estimated as *A_phys_* ≈ sin*φA_int_*. The maximum surface-to-volume ratio of the muscle is then:

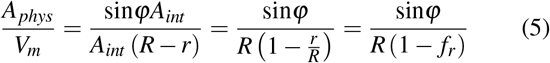

which is Eq. 2 in the main manuscript. The fraction of the available volume occupied by muscle is:

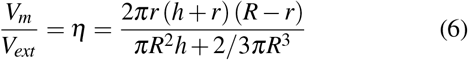

Introducing *r/R* = *f_r_* and *h/R* = *f_h_* yields Eq. 3 in the main manuscript:

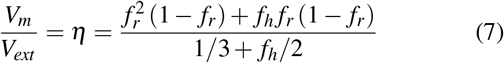

In the domain 0 < *f_r_* < 1 ^ *f_h_* ≤ 0, this function takes a maximum value:

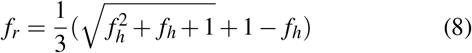

The value for *f_r_* which maximise *η* is thus at most 2/3 (for *f_h_* = 0), and then monotonously decays towards an asymptotic minimum value of *f_r_* = 0.5 as *f_h_* → ∞. These two limiting cases set an lower and upper bound for 0.44 < *η_max_* < 0.5, which can be shown formally be inserting Eq.8 in Eq. 7; the derivative of this function with respect to *f_h_* is positive for all *b* > 0. The two bounds for *η* may be understood by recognising that these limits correspond two ‘extreme’ shapes of a high-aspect ratio cylinder (*h* >> *r*) and a hemisphere (*h* = 0).

## Notes

### Competing Interest Statement

The authors have declared no competing interest.

